# The flagellar motor of *Vibrio alginolyticus* undergoes major structural remodeling during rotational switching

**DOI:** 10.1101/2020.04.24.060053

**Authors:** Brittany L. Carroll, Tatsuro Nishikino, Wangbiao Guo, Shiwei Zhu, Seiji Kojima, Michio Homma, Jun Liu

**Author notes:** Present address: Howard Hughes Medical Institute, Yale University School of Medicine, New Haven, Connecticut, USA. Both authors contributed equally to this work (Brittany L. Carroll, Tatsuro Nishikino). Author contributions: T.N., J.L. and M.H. designed research; S.Z., T.N., and S.K. performed experiments; B.L.C., T.N., J.L., W.G. and J.L. analyzed data; and B.L.C., T.N., J.L. and M.H. wrote the paper.

## Abstract

The bacterial flagellar motor is an intricate nanomachine that switches rotational directions between counterclockwise (CCW) and clockwise (CW) to direct the migration of the cell. The cytoplasmic ring (C-ring) of the motor, which is composed of FliG, FliM, and FliN, is known for controlling the rotational sense of the flagellum. However, the mechanism underlying rotational switching remains elusive. Here, we deployed cryo-electron tomography to visualize the C-ring in two rotational biased mutants (CCW-biased *fliG*-G214S and CW-locked *fliG*-G215A) in *Vibrio alginolyticus*. Sub-tomogram averaging was utilized to resolve two distinct conformations of the C-ring. Comparison of the C-ring structures in two rotational senses provide direct evidence that the C-ring undergoes major structural remodeling during rotational switch. Specifically, FliG conformational changes elicit a large rearrangement of the C-ring that coincides with rotational switching, whereas FliM and FliN form a spiral-shaped base of the C-ring, likely stabilizing the C-ring during the conformational remodeling.

## INTRODUCTION

Many bacteria navigate complex environments by controlling the flagellar rotational switch between counterclockwise (CCW) and clockwise (CW) in response to chemical stimuli. *Escherichia coli* and *Salmonella enterica* use a “run-and-tumble” approach for controlling movement, where flagella rotating in a CCW sense drives the cell body forward and when the rotation sense switches to CW the bacterium tumbles through the medium in an attempt to change direction (Berg, 2003; Chevance and Hughes, 2008; Terashima et al., 2008). *Vibrio alginolyticus* has a unique three step swimming pattern with forward, reverse, and flick motions; where CCW rotation propels the cell body forward, CW rotation drives the bacterium in reverse, and a flicking motion occurs upon CW-CCW rotation in an attempt to change swimming direction, analogous to the tumble (Xie et al., 2011).

The motor is the most intricate part of the flagellum that is not only responsible for flagellar assembly and rotation, but also essential for rotational switch. Spanning from the cytosol through the outer membrane, the motor consists of a series of rings, with the L-ring at the outer membrane, the P-ring located within the periplasmic space, the MS-ring embedded in the cytoplasmic membrane, and the C-ring inside the cytoplasm (Francis et al., 1992; Francis et al., 1994; Homma et al., 1987; Ueno et al., 1992). The stator subunits, embedded in the cytoplasmic membrane, generate torque via ion flow to rotate the C-ring (Blair, 2003; Sato and Homma, 2000b). Different from *E. coli* and *Salmonella*, *Vibrio alginolyticus* possesses several *Vibrio*-specific features: H, T, and O-rings. The O-ring is located on the outside of the outer membrane (Zhu et al., 2017), the H-ring facilitates the outer membrane penetration of the flagellum (Terashima et al., 2010; Zhu et al., 2018), and the T-ring contacts the H-ring and stators, presumably acting as a scaffold to hold the stators (Terashima et al., 2006; Zhu et al., 2019).

Flagellar rotation is powered by an electrochemical gradient across the cell membrane that drives ion flow through the stator complex (Berg, 2003; Kojima and Blair, 2004; Li et al., 2011; Terashima et al., 2008). In *Salmonella* or *E. coli*, H^+^ ions are conducted through the stator. In *Vibrio* species, Na^+^ ions are conducted through the stator. The prevailing idea is that MotA and MotB (in the H^+^ motor) or PomA and PomB (in the Na^+^ motor) form a membrane-bound stator subunit (Braun et al., 2004; Kojima and Blair, 2004; Sato and Homma, 2000a, b). The A and B subunits have four and one transmembrane helices, respectively, and two helices from the A subunit and one from the B subunit form a ion channel that contains an essential ion-binding aspartyl residue (Hosking et al., 2006). Inactive stators diffuse through the cytoplasmic membrane and interact with the rotor to generate a series of conformational changes within PomA and PomB that open of the ion channel and facilitate binding to the peptidoglycan layer (Fukuoka et al., 2009; Hosking et al., 2006; Kojima et al., 2018; Mino et al., 2019; Sudo et al., 2009; Zhu et al., 2014). However, exactly how the stator assembles around the motor and interacts with the C-ring is still not well understood.

The C-ring is essential for flagellar rotation and for changing the direction of rotation. The C-ring structure is conserved among diverse species, with repeating subunits consisting of FliG, FliM, and FliN, with FliN sometimes being supplemented or replaced by FliY (Kojima and Blair, 2004; Zhao et al., 1995). FliG, located at the top of the C-ring, interacts with the MS-ring, the lower C-ring, and the stator via three domains (Lee et al., 2010). The interaction of the N-terminal domain of FliG (FliG_N_) with FliF tethers the C-ring to the MS-ring and is necessary both for assembly and rotation of the flagellum (Lynch et al., 2017; Ogawa et al., 2015; Xue et al., 2018). The FliG_M_ domain interacts with FliM, holding the upper, membrane-proximal portion of the C-ring in contact with its lower, membrane-distal portion (Brown et al., 2002; Brown et al., 2007; Minamino et al., 2011). Lastly, the FliG_C_ domain interacts with a cytoplasmic loop of PomA (or MotA in proton-driven motors) via interactions of oppositely charged residues, connecting the stator and rotor (Lloyd and Blair, 1997; Takekawa et al., 2014; Yakushi et al., 2006). FliM also has three domains (Park et al., 2006). The N-terminal domain of FliM (FliM_N_) binds to the chemotaxis signaling protein phosphoryl CheY (CheY-P) to trigger switching from CCW to CW rotation (Paul et al., 2011; Vartanian et al., 2012). The FliM_M_ domain serves as a connection between the base of the C-ring and FliG (Brown et al., 2002; Minamino et al., 2011). FliM_C_ can form a heterodimer with FliN when fused via a flexible linker (Dos Santos et al., 2018; Notti et al., 2015). The third protein, FliN (and FliY in some species), a small single-domain protein, dimerizes with FliM or itself to form the base of the C-ring (Brown et al., 2005). Proposed models for FliG, FliM and FliN assembly include 1:1:4 (Sarkar et al., 2010a; Sarkar et al., 2010b) or a 1:1:3 (McDowell et al., 2016) stoichiometry. Two FliN homodimers form a ring at the base of each C-ring subunit in the first model (Sarkar et al., 2010a), while a FliM:FliN heterodimer and a FliN homodimer create a spiral base of the C-ring in the second model (McDowell et al., 2016).

It has been proposed that FliG undergoes a dramatic conformational change from an open, extended form during CCW rotation to a closed, compact form during CW rotation (Lee et al., 2010). Two α-helices play an important role in determining the FliG conformation: helix_MN_ connects FliG_N_ and FliG_M_, and helix_MC_ connects FliG_M_ and FliG_C_. These helices are extended in the open CCW conformation and become disordered, allowing for compaction, in the CW conformation. Each domain contains armadillo (ARM) repeat motifs; ARM_N_ interacts with the adjacent FliG monomer, and ARM_M_ and ARM_C_ interact either inter- or intramolecularly, depending upon the CCW or CW rotational sense, respectively (Brown et al., 2002; Lee et al., 2010; Minamino et al., 2011). This domain swapping polymerization is proposed to depend upon the rotational direction and to correspond with the conformational change in Helix_MC_ (Baker et al., 2016).

The FliG conformational change occurs about a hinge region first predicted by *in silico* model and characterized via mutational analysis (Van Way et al., 2004). Two recently characterized *fliG* variants in *V. alginolyticus* (G214S and G215A), previously characterized in *E. coli* (G194S and G195A (Van Way et al., 2004)), appear to hinder the conformational change sterically (Nishikino et al., 2018), resulting in motors that rotate primarily in a single direction. The *fliG*-G214S mutant produces a CCW-biased phenotype, whereas the *fliG*-G215A mutant produces a CW-locked phenotype. These adjacent residue substitutions create opposite motility phenotypes and are located in the Gly-Gly flexible linker, a hinge region whose conformation is believed to be dependent upon Helix_MC_ (Nishikino et al., 2016).

To characterize C-ring dynamics during switching, we used cryo-electron tomography (cryo-ET) to determine *in situ* motor structures of the *V. alginolyticus* CCW-biased mutant G214S and the CW-locked mutant G215A. We found that a large conformational change of the C-ring occurs between the CCW and CW motors. Docking the previously solved homologous structures into our cryo-ET maps, we for the first time build models of the C-ring in the conformations that correspond to CCW and CW rotation.

## RESULTS

### Visualization of intact flagellar motors in CCW and CW rotation states

To address the mechanism of rotational switching, we utilized cryo-ET to visualize the C-rings of on two FliG mutants (G214S and G215A) of *V. alginolyticus.* These were previously shown to have CCW-biased and CW-locked flagellar motors, respectively (Nishikino et al., 2018). Each *V. alginolyticus* cell usually possesses a single, polar, sheathed flagellum, which limits the amount of information per cell for *in situ* motor structure determination. We therefore introduced the *fliG*-G214S and *fliG*-G215A mutations (Table S1) into the *flhG* KK148 strain, which produces multiple flagella at the cell pole (Kusumoto et al., 2006). Multiple flagella can be readily seen in a typical cryo-ET reconstruction (Fig. 1A, E). In total, 2,221 CCW-biased motors from G214S cells and 1,618 CW-locked motors from G215A cells were used to determine *in situ* flagellar motor structures (Fig. 1). The structures derived from the CCW-biased and CW-locked mutants were similar in both size and shape (Fig. 1B, F). However, further refinement of the C-ring structures enabled us to resolve a 34-fold symmetry and two unique conformations of the C-ring (Fig. 1D, H).

**Figure 1.**
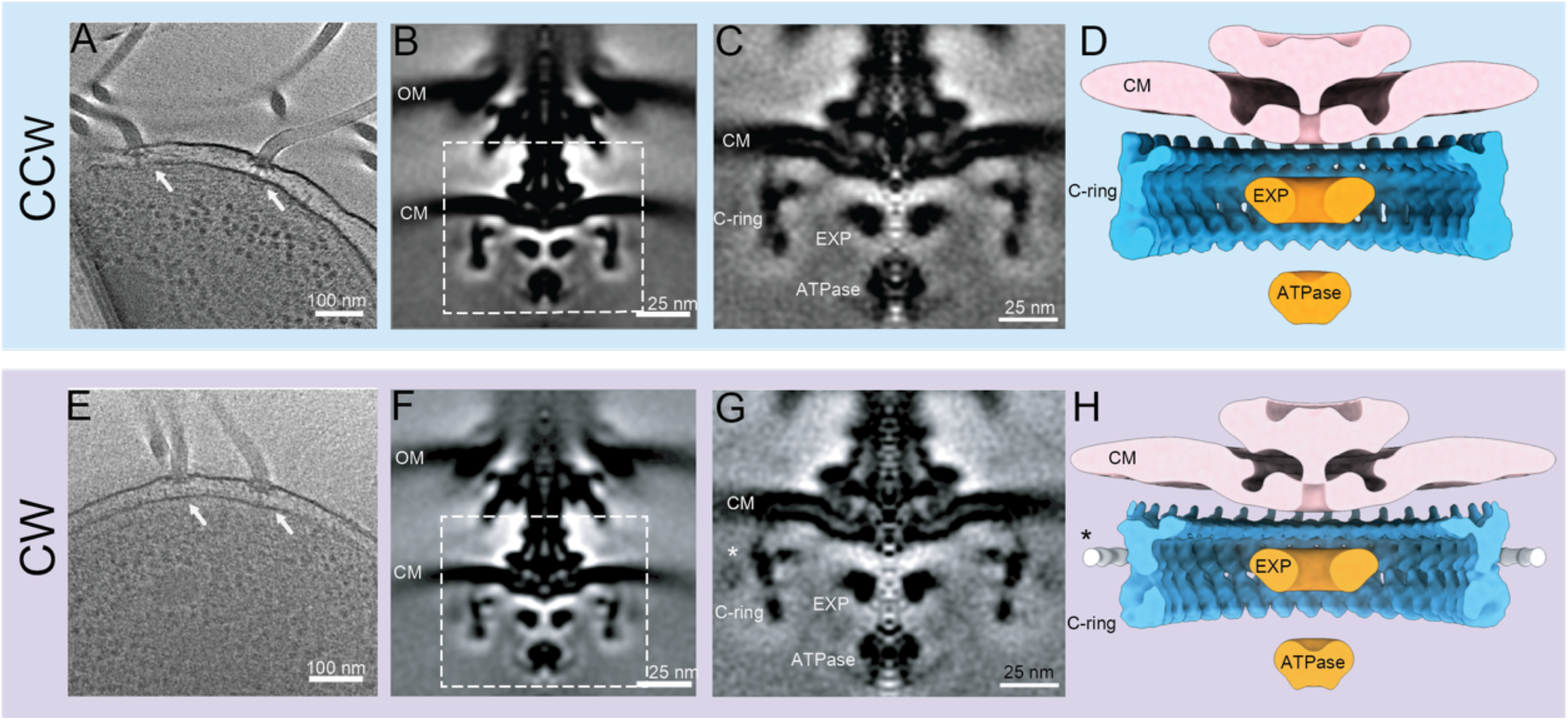
*In situ* structures of *V. alginolyticus* flagellar motor in CCW and CW rotation. (A) A representative tomogram slice showing two motors embedded in the cell envelope (white arrow, scale bar 100 nm). (B) The averaged structure of the CCW motor. (scale bar 25 nm). (C) The class average of the localized refinement of the C ring of the CCW motor (scale bar 25nm). (D) 3D reconstruction of the aligned CCW motor sliced down the middle of the X-axis in the same orientation as the class average. (E) A representative tomogram slice of the CW motor (white arrow, scale bar 100 nm). (F) The class average of the CW motor (scale bar 25 nm). (G) The class average of the localized refinement of the C ring of the CW motor (scale bar 25nm). (H) Half slice of the 3D reconstruction of the aligned CW motor. The asterisk represents additional density observed in only the CW motor. Abbreviations: outer membrane (OM), cytoplasmic membrane (CM), export apparatus (EXP)

It is evident from the 3D reconstructions that the C-ring undergoes a dramatic conformational change in switching rotational directions (Fig. 1D, H). There is a significant lateral tilt to the right upon switching from CCW to CW (Fig. 1D, H, S1). The overall diameter of the C-ring varies slightly between the two mutants. The diameter of the central portion of the C-ring in both mutants is 466Å. The bottom portion of C-ring has a spiral shape with a diameter of 490 Å in the CCW motor and 483Å in the CW motor. The top portion of the C-ring has a diameter of 466 Å in the CCW motor, while the diameter of the top portion of the CW motor is 486Å. Thus, the CW motor is about 20Å wider than the CCW motor at the top of the C-ring, even though the number of C-ring subunits remains 34 (Fig. 1, S1). These data demonstrate that the C-ring undergoes a large conformational change during flagellar switching despite experiencing limited changes in size and no change in composition of the C-ring.

### Molecular architectures of the CCW and CW C-rings

To understand the molecular detail, we built pseudoatomic models of the C-ring by docking available crystal structures from the C-ring proteins into the *in situ* maps derived from sub-tomogram averaging. First, we used homologous structures deposited in the PDB and SCWRL4 (Krivov et al., 2009) to map the *V. alginolyticus* amino acid sequence onto the open (3HJL; (Lee et al., 2010) and closed (4FHR; (Vartanian et al., 2012) conformations of the FliG structure. Second, we used I-TASSER (Roy et al., 2010; Yang et al., 2015) to generate the structures of FliM and FliN. Third, the models were placed into the cryo-ET map using UCSF Chimera (Pettersen et al., 2004). The known protein-protein interfaces were preserved during docking, and unknown protein-protein interfaces were refined with the Rosetta Protein-Protein docking function (Lyskov and Gray, 2008) before being placed into the cryo-ET map. Phenix (Afonine et al., 2013) was used to optimize the placement and to assess the overall fit using rigid body refinement.

We have modeled the C-ring with FliG at the top, FliM in the middle, and three FliN molecules at the base. Our model supports the 1:1:3 (FliG, FliM, and FliN) model in lieu of the 1:1:4 model, as there is no additional density for a fourth FliN molecule (Lee et al., 2010; Roy et al., 2010; Yang et al., 2015; Yang and Zhang, 2015). The 1:3 model suggests a FliM-FliN heterodimer interacts with the FliN homodimer characterized by Mass spectroscopy, as had been modeled into a previously solved cryo-EM map (McDowell et al., 2016; Thomas et al., 2006). We used this information and Rosetta Protein-Protein docking to identify and refine these interfaces in the *V. alginolyticus* flagellar motor. We modeled the CCW C-ring with a correlation coefficient (CC) mask (map-to-model) 0.74, CCbox 0.92 (Fig. 2). The CW C-ring had a CCmask of 0.68 and CCbox of 0.84 (Fig. 2).

**Figure 2.**
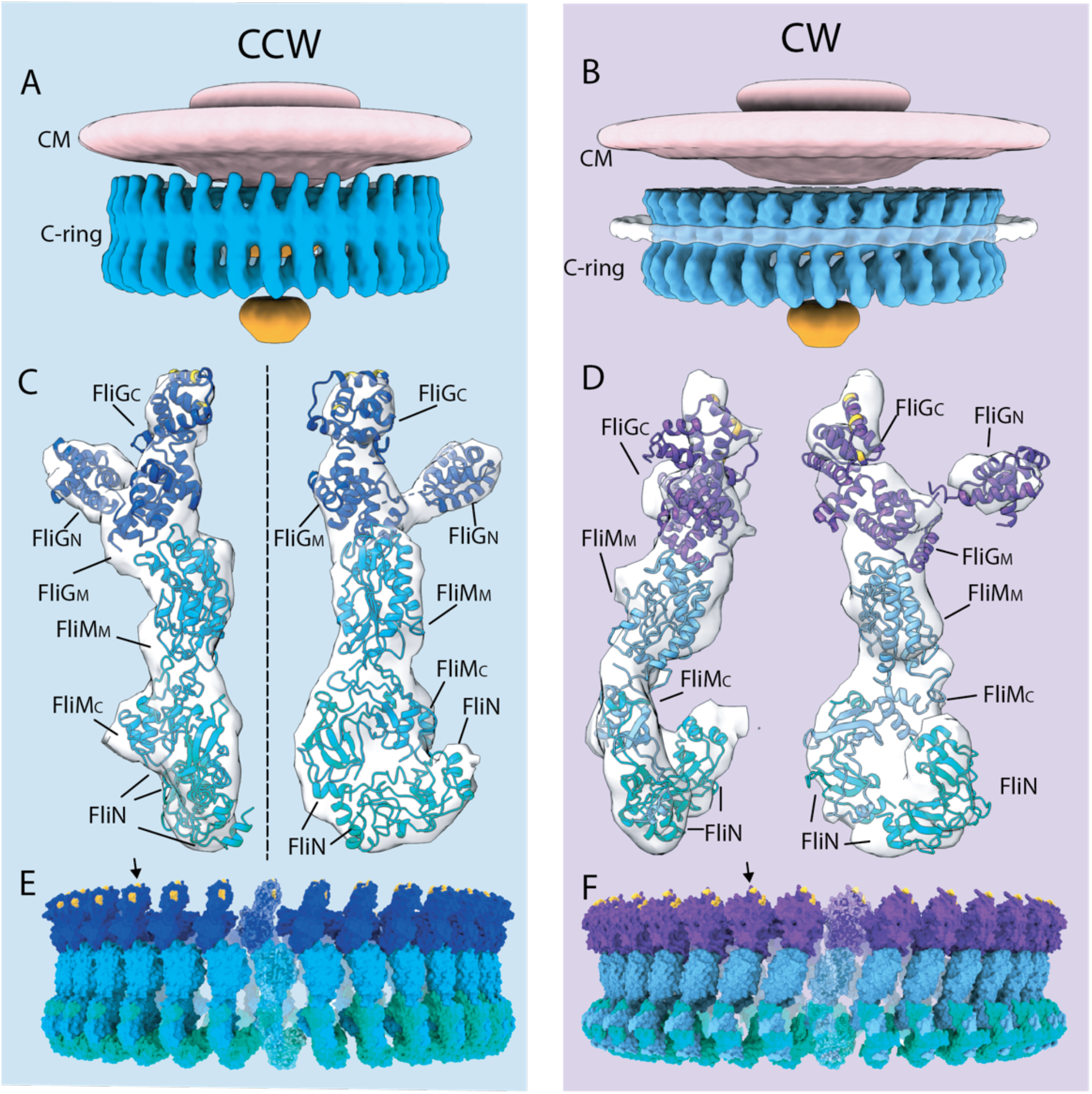
Models of the C-ring in CCW and CW rotation. 3D surface rendering of the CCW motor (A) and the CW motor (B). The MS-ring and cytoplasmic membrane (CM) is colored in pink, the C-ring in blue, and the export apparatus (EXP) in orange. (C)Two different views of the CCW-biased model of a single C-ring subunit fitted into the density (gray). FliG is shown in dark blue, FliM in light blue, and FliN in green. The charged residues that interact with the stator are shown in yellow. (D) The CW-locked model is shown as in (C), with FliG in purple, FliM in light blue, and FliN in green. The charged residues that interact with the stator are shown in yellow. (E) The CCW-biased model is expanded to 34-fold symmetry to complete the C-ring. (F) The CW-locked model is expanded to 34-fold symmetry creating the C-ring.

Our pseudoatomic models can be further substantiated from the biochemical data. The hydrophobic residues of ARM_C_ (Leu 216, Leu 219, Ile 222, Met 223, Leu 226, Leu 234, Ile 238, Met 245, and Ile 249) are poised to interact with ARM_M_ (Figure 2-figure supplement 1, orange sticks; (Park et al., 2006)). The EHPQR motif (Glu144, His145, Pro146, Gln147 and Arg179) of FliG is located in ARM_M_ and is orientated to interact with FliM_M_ (Figure 2-figure supplement 1, green sticks). Residues at the FliM-FliM interface had previously been identified in *Salmonella* and *E. coli* correspond to Asn56, His63 Asp184, Pro185, and Met187 in *V. alginolyticus* (Park et al., 2006; Sakai et al., 2019). The N-terminus of FliM_M_ is orientated facing outward toward the additional density of the CW-locked mutant in a position in which that it could bind to CheY-P (Figure 2-figure supplement 1, asterisk and E) (Lee et al., 2001). Our models of the C-ring are thus consistent with both the previous biochemical data and our cryo-ET maps.

### FliM/FliN interactions hold the C-ring subunits together

The tilting observed between the CCW and CW motors is centered on FliM, which serves as a structural protein that holds the C-ring subunits together. However, without a full-length crystal structure, the relative orientations of FliM_N_, FliM_M_ and FliM_C_ are unknown. The previous crystal structures and solution NMR studies showed FliM_M_ interacting with FliG (Dyer et al., 2009; Lam et al., 2013; Vartanian et al., 2012). FliM_C_ shows sequence homology to FliN and structural homology to the FliN homodimer, as shown from the structure of a crystallized FliM_C_-FliN fusion protein (McDowell et al., 2016; Notti et al., 2015). To place FliM in our structure we generated a FliM_MC_ model using I-TASSER and oriented the domain orientation to fit into our cryo-ET map (Fig. 3). We lacked sufficient information to confidently place FliM_N_, and we therefore left it out of our model. The position of FliM_N_, which binds CheY-P, is flexible (discussed below). Our model shows that FliM_M_ forms the middle portion of the C-ring and allows for the tilting of the C-ring, as FliM_C_ tethers FliM to the spiral base via extensive contacts with FliN in the heterodimer. The spiral base is largely built by the smallest protein in the C-ring, FliN, with alternating FliM-FliN heterodimers and FliN homodimers (Fig. 3). The spiral itself remains largely unaltered upon rotational switching (Fig. 3 & S1). The rigidity of the spiral suggests that it has the important function of keeping the C-ring subunits connected during the conformational remodeling.

**Figure 3.**
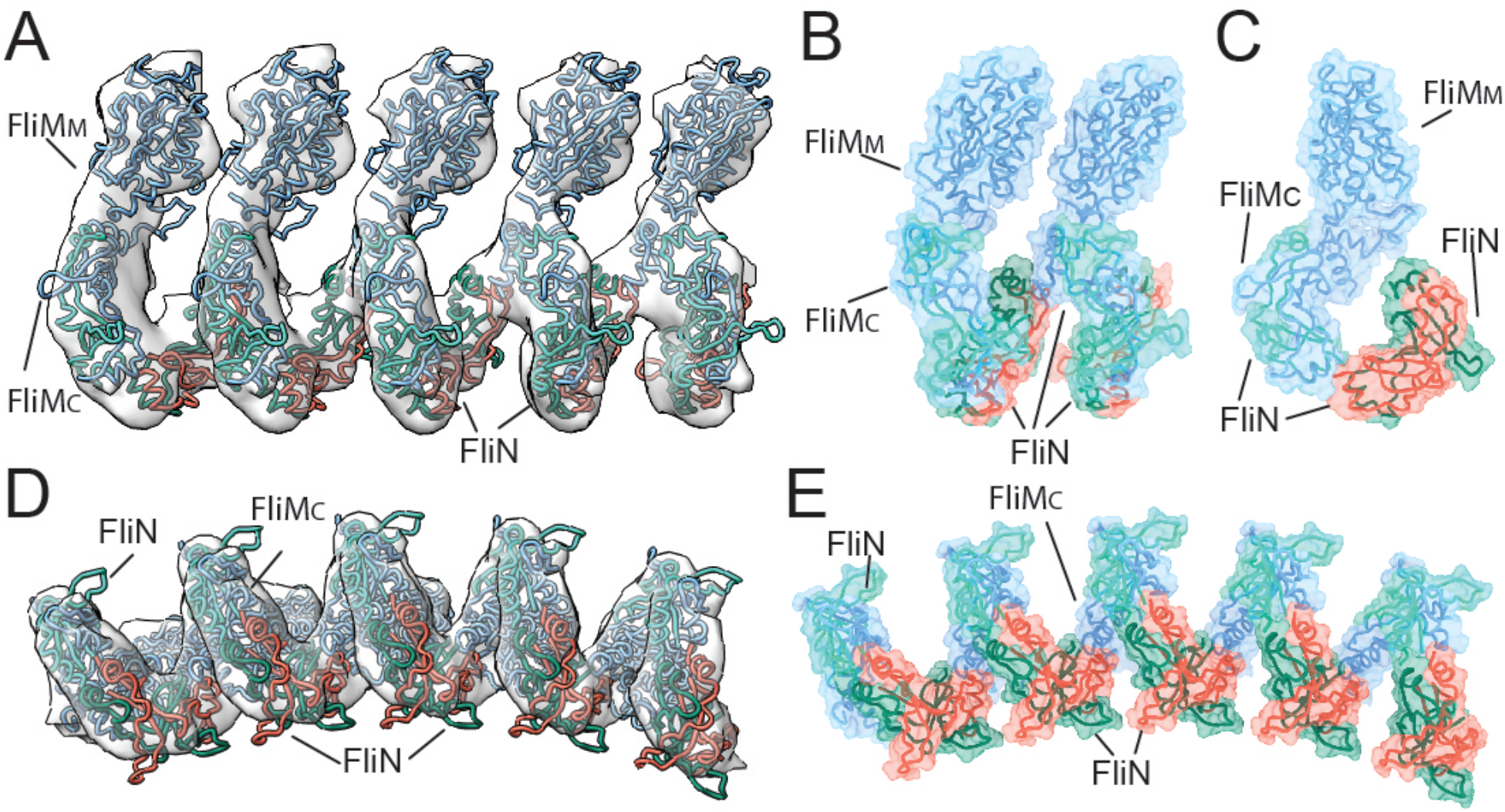
FliM_C_ and FliN form a continuous spiral at the base of the C-ring. (A) Five C-ring subunits of FliM and FliN are shown as a licorice fit into the cryo-ET map (white). FliM is shown in blue, and the three individual FliN molecules are shown in green, orange, and dark green. (B) Two C-ring subunits are depicted as licorice, with surfaces shown. (C) A single C-ring subunit, represented as licorice and surfaces, rotated 90° to show the spiral from the side looking down the hole. View from the bottom of the C-ring showing the spiral in the (D) density map and (E) represented as surfaces.

### Presentation of FliGc to the stator in the CCW and CW motors

To determine which proteins were responsible for the tilting we observed, we expanded our subunit model to the whole C-ring model by applying 34-fold symmetry (Fig. 4). The expansion of the model preserved the spiral at the base observed in the maps, highlighting the similarities of the base in the CCW and CW motors (Figure 1-figure supplement 1). A superposition of the CCW and CW C-ring subunits in the CCW and CW motors revealed a 31Å shift of FliG with a 36° tilt about FliM (Fig. 4A). FliG and FliM tilt 25° outward upon switching from CCW to CW rotation (Fig. 4B). The movements of FliG and FliM result in a 89° rotation of the charged residues through a movement of between 35 to 40 Å (Fig. 4C). This suggests that, upon switching, FliG is presumably presented to the stator units very differently as a cause or as a result of lateral tilting of C-ring.

**Figure 4.**
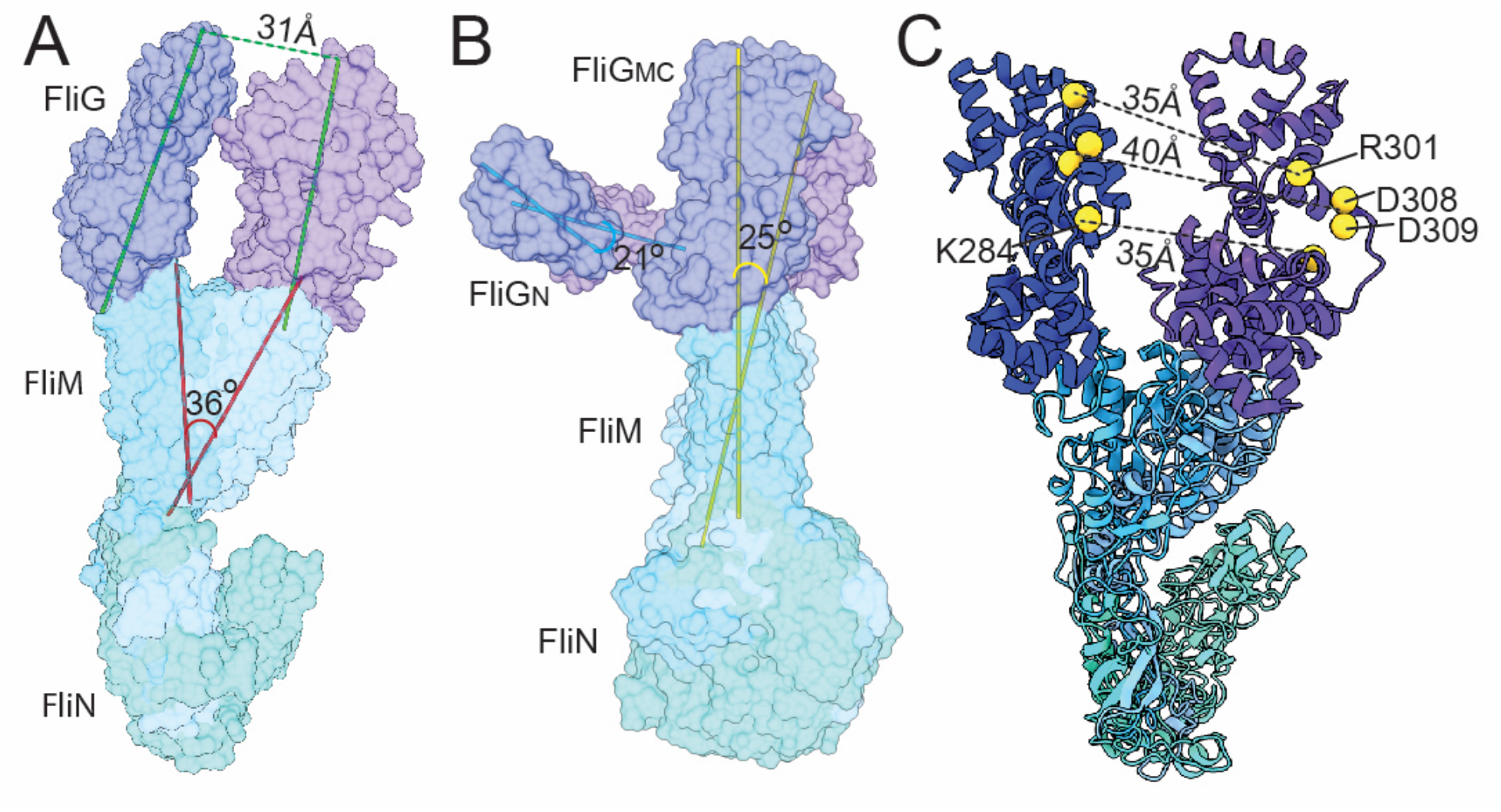
The FliG conformational change results in a tilt about FliM to change the presentation of FliG_C_ to the stator. (A) The CCW-biased and CW-locked models at the same position of the C-ring are shown from the front. The CCW-biased model with FliG (dark blue), FliM (light blue), and FliN (green); the CW-locked model with FliG (purple), FliM (light blue), and FliN (green). The axes for the center of mass of FliG_MC_ are shown in green, those for FliM_M_ are shown in red. The FliG subunits move approximately 31Å with a 36° tilt of FliM_M_ laterally upon switching, as seen in the front and back views. (B) The same models are rotated 90° to show the comparison of the C-ring subunit from the side. The axes for the center of mass of FliG_MC_/FliM is shown in yellow, and those for FliG_N_ are shown in blue. The conformational change of the C-ring results in an approximately 25° tilt of mass outward upon switching from CCW to CW rotation. The spiral at the base of FliN remains stationary relative to FliM_M_ and FliG. (C) The charged residues that interact with the stator rotate 89°, resulting in 35-40 Å difference in position.

### Extra diffuse density around the CW C-ring

We observed a density around the CW-locked C-ring that is most likely CheY-P (Fig. 1). Although the resolution is too low to verify that this density is formed by CheY-P, we have several reasons to believe that it is. First, the density is above the noise level in the CW motors but not in the CCW motors. Second, Chang et al. (in preparation) have used GFP-tagged CheY-P to show that, in *Borrelia burgdorferi*, CheY-P occupies the similar position. Third, when we place the crystal structure of CheY into the model, there is about 60Å gap between the C-ring density and this additional density (Figure 2-figure supplement 1). This distance could be bridged by the 42 residues of FliM_N_ to connect CheY-P to FliM_M_. The density may be diffuse as there may every FliM_N_ may not be bound by CheY-P.

## DISCUSSION

The flagellar motor structures we determined provide direct evidence that the CCW and CW forms possess two distinct conformations of the C-ring. We observed a 36° lateral and 25° medial tilting of the C-ring, which almost certainly alter the interactions between C-ring and the stator. The exact position of the stator and how it interacts with C-ring in *Vibrio* is unknown, as the cytoplasmic portion of the stator has yet to be visualized by cryo-ET. In our pseudoatomic model, the charged residues (Lys284, Arg301, Asp308, and Asp309 in FliG_C_) that are known to interact with the stator are located at the top of the C-ring and move about 35 Å with an 85° rotation. This movement of the charged residues is plausible as a flexible region connects FliG_CC_ (where the charged residues are located) and FliG_CN._ This movement is possibly necessary to create critical interactions between the stator and the C-ring in the two conformations. FliG_CN_ contains the ARM_C_ motif that interacts with FliG_M_ and FliM_M_, and FliG_CC_ contains the charged residues that interact with the stator. The flexible linker allows for a large range of movement in FliG_CC_ relative to FliG_CN_ and FliG_M_, thus suggesting that the presentation of the charged residues could vary greatly depending upon the conformation of FliG. The dynamics of the C-ring appear to be confined to the upper two-thirds of the structure, with the spiral base that connects the C-ring subunits remaining stable.

Notably, our cryo-ET model shows 34-fold symmetry for both the CCW and CW rotating C-rings, with only a modest diameter change in FliG. It has previously been hypothesized, based on experiments utilizing fluorescently tagged FliM and FliN, that the FliG, FliM and FliN composition of the C-ring changes during rotational switching (Delalez et al., 2014; Delalez et al., 2010; Hosu and Berg, 2018; Lele et al., 2012). A recent single-particle cryo-EM study suggested that inter-subunit spacing and interactions of the C-ring account for motor switching in *Salmonella* (Sakai et al., 2019). That study reported a 9Å decrease in C-ring diameter upon switching to CW, opposite to what we observed, a 20Å increase in the diameter at FliG_C_ in the C-ring of the CW *V. alginolyticus* motor. These differences are modest and could simply be due to differences in the bacterial species or to the differences in the techniques employed. Whatever the explanation for the discrepancies may be, our cryo-ET data suggest that the composition of the C-ring remains constant and that the change in rotational direction occurs from changes in the presentation of the charged residues of FliG_cc_ to the stator.

Our cryo-ET maps confirm that FliG, FliM and FliN interact in a 1:1:3 ratios, as suggested for the non-flagellar homolog Spa33 in *Shigella* (McDowell et al., 2016). This stoichiometry favors a model of the C-ring in which two homodimers of FliN creating a closed loop. By modeling the complete structure of the C-ring, we have shown that FliM holds the individual C-ring subunits together by interacting with FliG_M_ and forming a heterodimer with FliN, anchoring the stalk into the spiral. The FliM-FliN spiral connects adjacent C-ring subunits, allowing for dramatic movement without dissociation of the subunits. The FliG_M_–FliM_M_ interface, perhaps dynamic in its own right, has been suggested to be involved in flagellar switching (Dyer et al., 2009; Pandini et al., 2016; Sakai et al., 2019). It was shown, using NMR, that the CheY-P binding to FliM displaces FliG_C_, and that the dramatic rearrangement of FliG_MC_ is possible because of the flexibility of FliG (Dyer et al., 2009). In particular, the GGPG loop in FliM_M_ is suggested to be critical for rotation of FliG_M_ relative to FliM_M_ (Pandini et al., 2016). Most recently, point mutations targeting FliM_M_ were shown to restore CCW rotation in a CW-biased mutant (Sakai et al., 2019). These results, combined with our findings, lead us to suggest that the conformational change in FliG results in a rotation about FliM_M_ that leads to the tilting of the C-ring subunit to alter the presentation of the charged residues of FliG_cc_ to the stator.

We propose a model for CCW-to-CW switching initiated by CheY-P binding (Fig. 5). CheY-P binding triggers a tilting about FliG and FliM, that results in a conformational change in FliG and alters the interactions between the C-ring and stator. FliM_C_ and FliN create a spiral base that holds the C-ring together during these dynamic rearrangements. FliM_C_ and FliN may also be involved in relaying the conformational change to adjacent C-ring subunits. By placing the FliG crystal structures of the open and closed variations within our cryo-ET image, we provide additional evidence to support the proposed model where the ARM_M_ and ARM_C_ domains toggle between inter- and intramolecular interactions (Lee et al., 2010). NMR data suggest that FliG_C_ is the domain that moves and that FliG_M_ remains in stable contact with FliM (Dyer et al., 2009). These changes in the C-ring structure produce the two directions of flagellar rotation.

**Figure 5.**
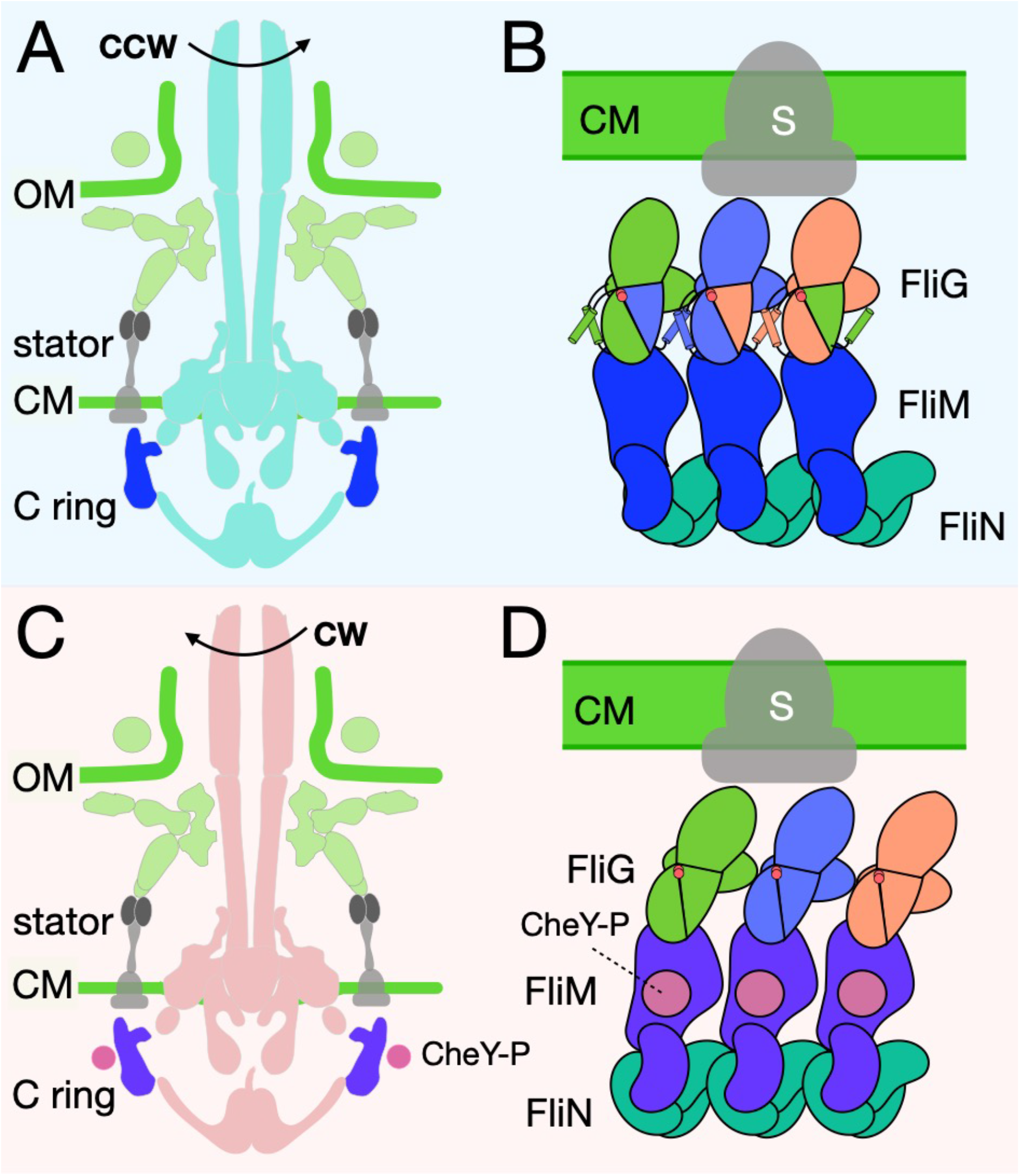
A model for the switching mechanism. (A) A cartoon of the intact motor with the previously resolved structures in light blue or light purple, the C-ring in blue or purple, the unresolved portion of the stator, PomA, in grey, and CheY-P in pink. (B) A cartoon representation of the C-ring rearrangement upon CheY-P binding and rotational switching to the CW sense. In CCW rotation, the intermolecular interactions of FliG ARM_M_ and ARM_C_ are possible because helix_NM_ and helix_MC_ are ordered. The Gly-Gly flexible region is depicted by red circles. The stator (gray) is shown to interact with FiG_c_. Upon CheY-P binding, the C-ring undergoes a conformational change that produces CW rotation. During this transition, the center of mass tilts laterally and slightly outward. In CW rotation there are intramolecular interactions of FliG ARM_M_ and ARM_C_. CheY-P (dark purple) is shown bound to FliM, and the stator (gray) interacts with FliG_C._ We hypothesize that the conformational change in FliG is initiated *in vivo* by CheY-P binding, and this switches the rotational direction of the C-ring by changing how FliG_C_ interacts with the stator. FliG undergoes a conformational change and FliM tilts about the base of the spiral. The spiral keeps the C-ring subunits connected during these large changes as the interactions within the FliG ARM domains toggle between inter- and intramolecular.

Understanding the interactions of the C-ring with the stator and the MS-ring is essential to elucidate the mechanism of rotational switching and transmission of the rotation to the flagellar filament. We could not resolve the cytoplasmic portion of the stator in our cryo-ET map, and thus we could not confirm directly that the orientation of FliG_CC_ alters the presentation of its charged residues to the charged residues of PomA. Restricting the movement of FliG_CC_ via truncations of the flexible linker may address the importance of FliG_CC_ mobility.

The interaction of FliG_N_ with FliF of the MS-ring is necessary for assembly of the flagellar motor. A recent cryo-EM structure shows that various isoforms of FliF oligomerize to form the MS-ring, presenting 34-fold symmetry to the C-ring and thus explaining the long-standing mismatch issue (Johnson et al., 2019). Visualization of the FliG_N_ – FliF interface in both the CCW and CW models will elucidate whether this interaction is static or dynamic. With the rapid development of cryo-ET, it has potential to reveal motor structure *in situ* at higher resolution, which will further our understanding of the flagellar assembly and function.

In summary, we have shown a major conformational change of the C-ring due to a single point mutation in FliG. Using the cryo-ET map, we generated a molecular model that attributes the tilt component of the conformational change to FliM. This phenomenon suggests rotation of FliG about FliM presents the charged residues of FliG_CC_ to the stator in a manner that controls the rotational sense. These movements within the C-ring subunits may be relayed throughout the switch complex by interactions between FliM and FliN within the spiral. We propose that the switching complex is dynamically repositioned depending upon the rotational direction in a way that is governed by CheY-P association with and dissociation from FliM. By manipulating the protein expression of CheA and CheZ, the enzymes responsible for controlling CheY phosphorylation we will be able to elucidate the role of CheY during switching at the molecular level. To understand how torque is generated during CCW and CW rotation, a direct visualization of the rotor – stator interaction will be necessary.

## MATERIALS AND METHODS

### Bacterial strains, plasmids, and growth condition

Bacterial strains used in this study are listed in Table S1. To introduce the *fliG* deletion, NMB328 was constructed from KK148 using pHIDA3 by allelic exchange as previous reported (Kusumoto et al., 2006; Le Roux et al., 2007). *V. alginolyticus* strains were cultured at 30°C on VC medium (0.5% [wt/vol] polypeptone, 0.5% [wt/vol] yeast extract, 3% [wt/vol] NaCl, 0.4% [wt/vol] K_2_HPO_4_, 0.2% [wt/vol] glucose) or VPG medium (1%[wt/vol] polypeptone, 3% [wt/vol] NaCl, 0.4% [wt/vol] K_2_HPO_4_, 0.5% [wt/vol] glycerol). If needed, chloramphenicol and Isopropyl β-D-1-thiogalactopyranoside (IPTG) were added at final concentrations of 2.5 µg/ml and 1 mM, respectively. *E. coli* was cultured at 37°C in LB medium (1% [wt/vol] Bacto tryptone, 0.5% [wt/vol] yeast extract, 0.5% [wt/vol] NaCl). When culturing *E. coli* β3914 strain, 2,6-diaminopimelic acid was added to the LB medium to a final concentration of 300 µM. If needed, chloramphenicol was added at final concentrations of 25 µg/ml.

### Mutagenesis

To introduce mutations (G214S or G215A) in the *fliG* gene on plasmid pNT1 site-directed mutagenesis was performed using the QuikChange method, as described by the manufacturer (Stratagene). All constructs were confirmed by DNA sequencing. Transformation of *V. alginolyticus* with plasmid pNT1 was performed by conjugational transfer from *E. coli* S17-1, as described previously (Okunishi et al., 1996).

### Sample Preparation

The methods of sample preparation, data collection, data processing and sub-tomogram analysis were followed as described previously (Zhu et al., 2018). *V. alginolyticus* cells were cultured overnight at 30°C on VC medium and diluted 100 X with fresh VPG medium and cultured at 30°C. After 4 or 5 hours, cells were collected and washed twice and finally diluted with TMN500 medium (50 mM Tris-HCl at pH 7.5, 5 mM glucose, 5 mM MgCl, and 500 mM NaCl). Colloidal gold solution (10-nm diameter) was added to the diluted *Vibrio* sp. samples to yield a 10 X dilution and then deposited on a freshly glow-discharged, holey carbon grid for 1 min. The grid was blotted with filter paper and rapidly plunge-frozen in liquid e.

### Data collection and processing

The frozen-hydrated specimens of NMB328 was transferred to a Titan Krios electron microscope (Thermo Fisher Scientific). The microscopes are equipped with a 300-kV field emission gun (FEG) and a direct electron detector (Gatan K2 Summit). The images were collected at a defocus near to 0 µm using a Volta phase plate and the energy filter with a 20-eV slit. The data were acquired automatically with SerialEM software (Mastronarde, 2005). For better data collection, the phase shift is normally distributed in the range of 0.33π to 0.67π. A total dose of 50 e^−^/Å^2^ was distributed among 35 tilt images covering angles from −51° to +51° at tilt steps of 3°. For every single tilt series collection, the dose-fractionated mode was used to generate 8 to 10 frames per projection image. Collected dose-fractionated data were first subjected to the motion correction program to generate drift-corrected stack files (Li et al., 2013; Morado et al., 2016). The stack files were aligned using gold fiducial markers and volumes reconstructed using IMOD and Tomo3d, respectively (Agulleiro and Fernandez, 2015; Kremer et al., 1996). In total, 259 tomograms of CCW state motor (G214S mutation) and 220 tomograms of CW-state motor (G215A mutation) were generated.

### Subtomogram analysis with I3 package

Bacterial flagellar motors were detected manually, using the I3 program (Winkler, 2007; Winkler et al., 2009). We selected two points on each motor, one point at the C-ring region and another near the flagellar hook. The orientation and geographic coordinates of selected particles were estimated. In total, 1,618 and 2,221 sub-tomograms of *V. alginolyticus* motors from CW-motor and CCW-motor, respectively, were used for sub-tomogram analysis. The I3 tomographic package was used on the basis of the “alignment by classification” method with missing wedge compensation for generating the averaged structure of the motor, as described previously (Zhu et al., 2017).

### Model Generation

The *V. alginolyticus* C-ring proteins were generated using I-TASSER (Roy et al., 2010; Yang et al., 2015). The proteins were trimmed to the homologous structures deposited in the PDB to avoid large clashes. Using I-TASSER we have generated two models of *Vibrio* FliM, FliM full length, and FliM_C_ with the linker region before. The full length FliM generated by I-TASSER had the correct topology for FliM_M_, but the orientation FliM_C_ relative to FliM_M_ was incorrect. Furthermore, the folding of FliM_C_ is similar to the previously solved crystal structure, but different enough that the FliM-FliN heterodimer was unable to be modeled. To circumvent this problem, we ran I-TASSER with just the last 223 residues of FliM, including the FliM_C_ and the flexible region immediately before. Due to the unknown relative location of the flexible regions it was necessary to trim. FliG, FliM, and FliN were placed into the CCW-biased and CW-locked cryo-ET maps by hand guided by the literature, and fit using the Chimera fit to map function. Mainly, four PDB models were used the full length FliG (3HJL; (Lee et al., 2010)), FliG-FliM (4FHR; (Vartanian et al., 2012)), FliM-FliN fusion (4YXB; (Notti et al., 2015)), and the FliN-dimer (1YAB; (Brown et al., 2005)).

### Model refinement

The model was refined using PHENIX Real Space Refinement (Afonine et al., 2013) to move the protein domains relative to one another while preserving the known architecture of the C-ring subunits. The unknown protein-protein interfaces were refined in Rosetta using the protein-protein docking scripts (Lyskov and Gray, 2008). This optimized protein model was then rigid body refined using PHENIX Real Space Refinement.

## ACKNOWLEDGEMENTS

We thank Michael Manson for critical reading and suggestions. We thank A. Abe in our laboratory for technical support in this research. This work was supported in part by JSPS KAKENHI Grant Numbers JP16H04774 and JP18K19293 (to S.K.), JP18K06155 (to T.K), and Program for leading Graduate Schools of Japan, Science for the Promotion of Science (JP17J11237 to T.N.). T.N. was supported in part by the Integrative Graduate Education and Research program of Nagoya University. B.L.C., S.Z. and J.L. were supported by grants GM107629 and R01AI087946 from National Institutes of Health.

## SUPPORTING INFORMATION

Supplementary information associated with this article can be found in the online version, at the publisher’s website.

## SUPPLEMENTAL INFORMATION

**Table S1.**
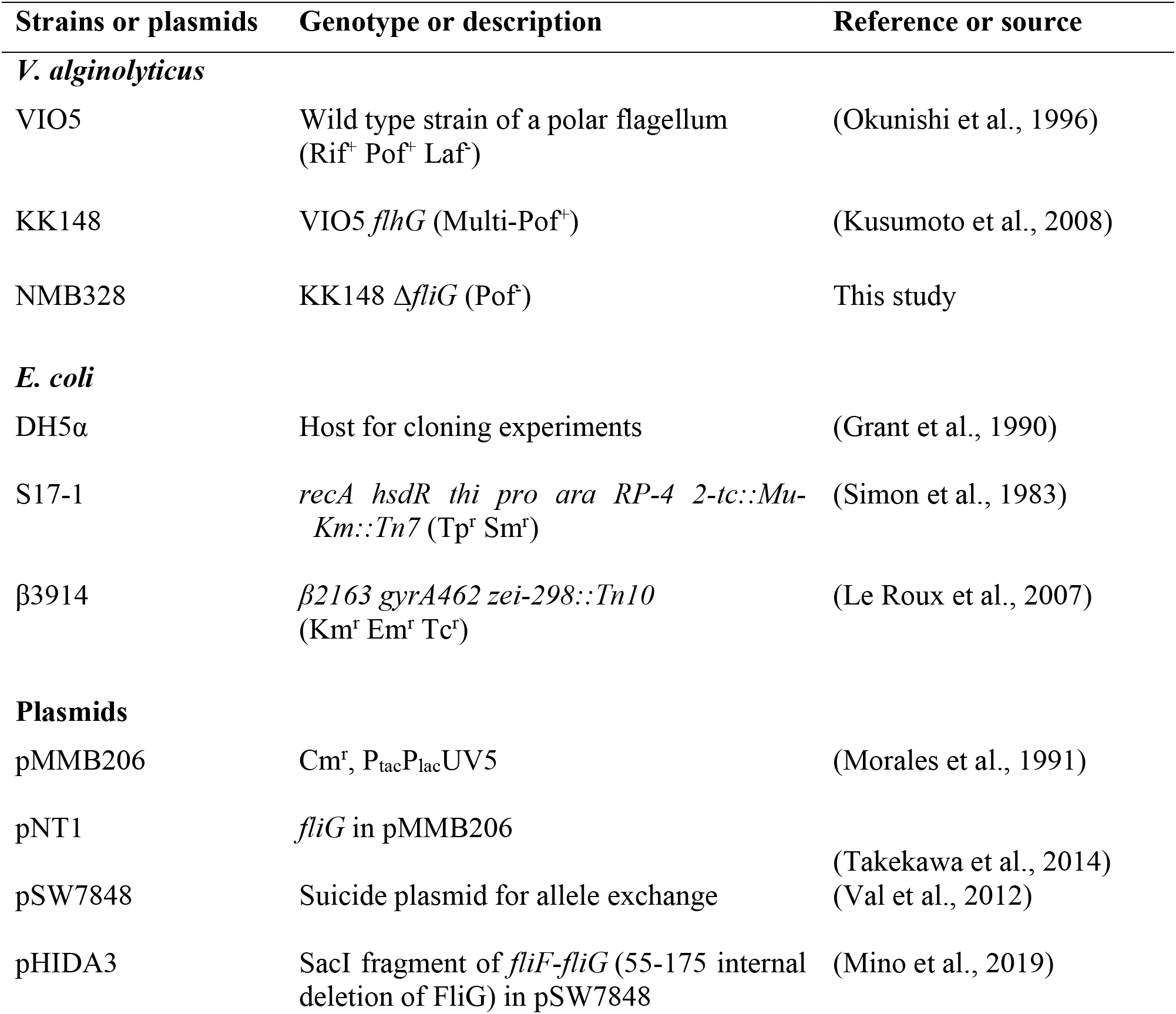
Strains and plasmids.

**Table S2:** Cryo-ET data used in this study.

| Strain genotype | Tomograms | Motors |
| --- | --- | --- |
| *fliG-G214S* | 259 | 2,221 |
| *fliG-G215A* | 220 | 1,618 |

**Table S3:** The C-ring diameter comparison between CCW and CW rotations.

|  | Top | Middle | Bottom |
| --- | --- | --- | --- |
| CCW | 466 Å | 466 Å | 490 Å |
| CW | 486 Å | 466 Å | 483 Å |

**Figure 1-figure supplement 1.**
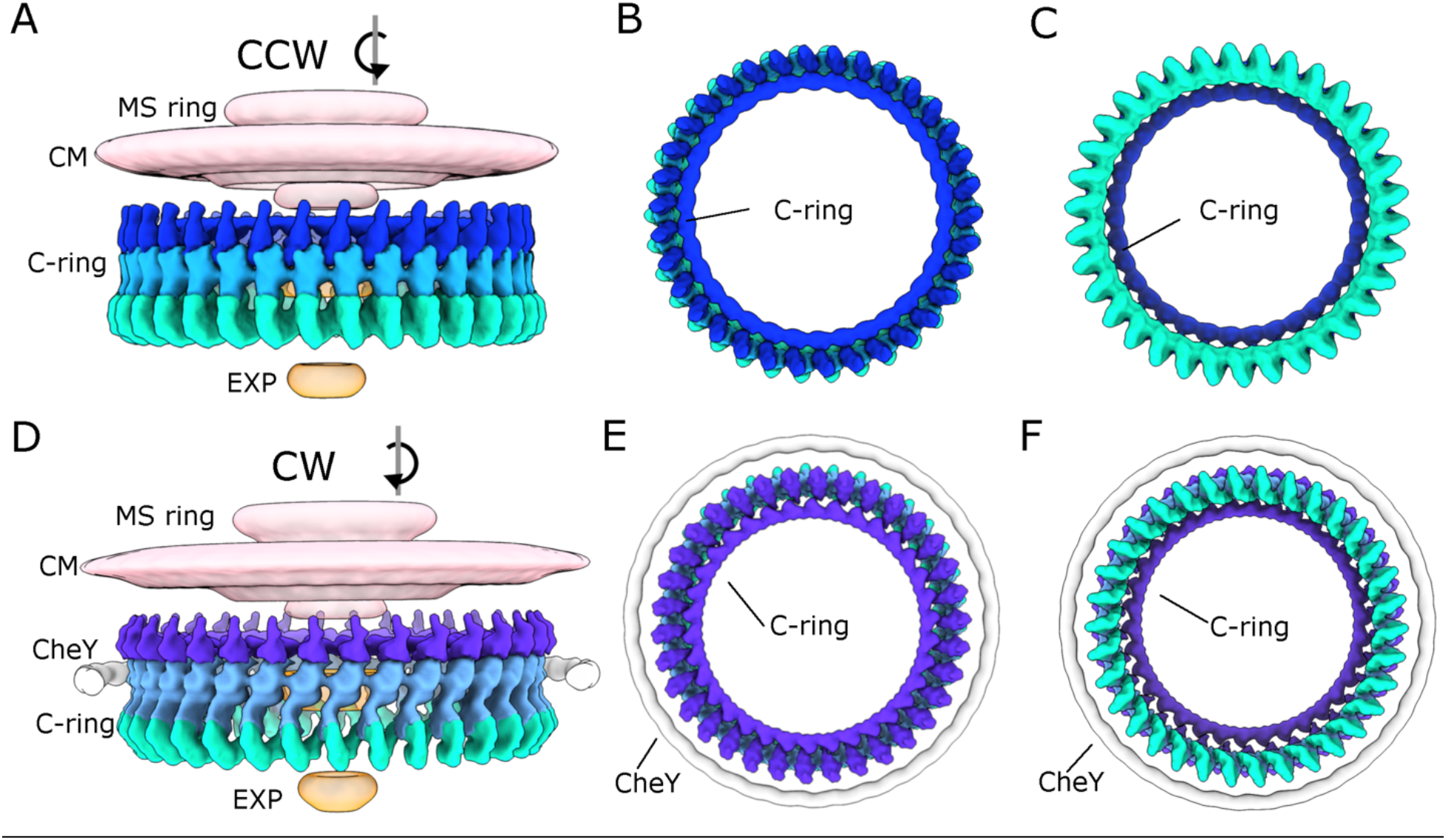
The switch complex tilts ~36° between the CCW to CW rotation. (A) A side view of the cryo-ET map of the C-ring in CCW rotation. The MS-ring and cytoplasmic membrane (CM) is colored in pink, export apparatus in orange. The C-ring is further segmented into FliG (blue), FliM (light blue), and FliN (green). (B and C) A top view and a bottom view of the C-ring in CCW rotation, respectively. (D) A cryo-ET map of the CW C-ring is segmented as CM and MS-ring (pink), EXP (orange), CheY (grey), and the C-ring proteins FliG (purple), FliM (grey-blue) and FliN (green). (E and F) A top and bottom view, respectively. The connectivity of the spiral formed by FliM_C_ and FliN can be seen clearly in the side and bottom views.

**Figure 2-figure supplement 1.**
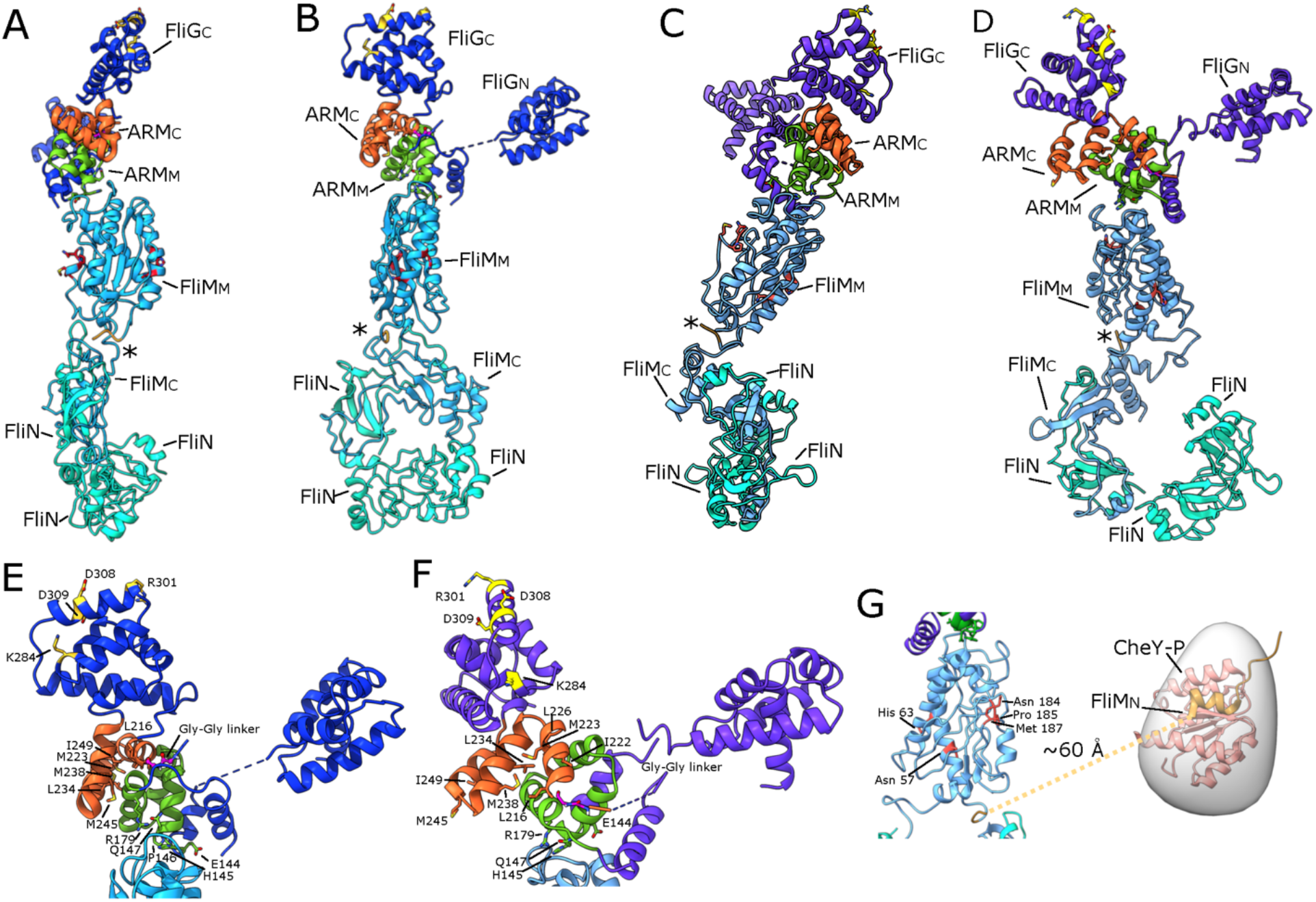
The important regions biochemically and structurally of the C-ring in the CCW and CW models. Previous biochemical and structural work has elucidated many important residues of the C-ring subunits. Here we have highlighted these regions in our complete model. (A) The CCW-biased model is shown from the front, with FliG in blue, ARM_M_ in green, ARM_C_ in orange, FliM in blue, and FliN in teal. (B) The CCW-biased model from the side. (C) The CW-locked model, FliG in purple, ARM_M_ in green, ARM_C_ in orange, FliM in blue, and FliN in teal from front (D) side view. The red residues in FliM_M_ are residues that have been shown to interact with neighboring FliM molecules in *E. coli* or *Salmonella*. The residues in our model are located on the sides of FliM such that they would be able to interact with a neighboring C-ring subunit. FliM is orientated such that the N-terminus of FliM (tan, asterisk) would be able to extend from FliM to interact with CheY-P. A close up on FliG in CCW-biased (E) and CW-locked (F) shows the charged residues Lys 284, Arg 301, Asp 308, and Asp 309 (yellow sticks) that have been shown to interact with the stator (Nishikino et al., 2016). The important hydrophobic residues, Leu 216, Ala 219, Ile 222, Met 223, Leu 226, Leu 234, Met 238, Met 245, and Ile 249 (orange sticks (Park et al., 2006)), and the EQPHR residues (green sticks) show the interactions of FliG_MC_ and FliM_M_ (Park et al., 2006; Sakai et al., 2019). (G) The CW-locked FliM_M_, is depicted in cartoon with CheY-P (1F4V, (Lee et al., 2001)) placed in the the extra density around the C-ring (white). This shows that FliM could interact with CheY-P if the extra density is in fact CheY-P.

